# Concerted regulation of skeletal muscle metabolism and contractile properties by the orphan nuclear receptor Nr2f6

**DOI:** 10.1101/2023.05.17.540900

**Authors:** Dimitrius Santiago P.S.F. Guimarães, Ninon M.F. Barrios, André Gustavo de Oliveira, David Rizo-Roca, Maxence Jollet, Jonathon A.B. Smith, Thiago R. Araujo, Marcos Vinicius da Cruz, Emilio Marconato Junior, Sandro M. Hirabara, André S. Vieira, Anna Krook, Juleen R. Zierath, Leonardo R. Silveira

## Abstract

**Objective:** The maintenance of skeletal muscle plasticity upon changes in the environment, nutrient supply, and exercise depends on regulatory mechanisms that couple structural and metabolic adaptations. However, the mechanisms that interconnect both processes at the transcriptional level remain underexplored. Nr2f6, a nuclear receptor, regulates metabolism and cell differentiation in peripheral tissues. However, its role in the skeletal muscle is still elusive. Here, we aimed to investigate, for the first time, the effects of Nr2f6 modulation on muscle biology *in vivo* and *in vitro*.

**Methods:** Global RNA-seq was performed in Nr2f6-knockdown C2C12 myocytes (N=4-5). Molecular and metabolic assays and proliferation experiments were performed using stable Nr2f6 knockdown and overexpression C2C12 cell lines (N=3-6). Nr2f6 content was evaluated in *in vitro* and *in vivo* lipid overload models (N=3-6). *In vivo* experiments included Nr2f6 overexpression in mouse tibialis anterior muscle, followed by gene array transcriptomics and molecular assays (N=4), *ex vivo* contractility experiments (N=5), and histological analysis (N=7). The conservation of Nr2f6 depletion effects was confirmed in primary human and mouse skeletal muscle cells.

**Results:** Nr2f6 knockdown upregulated genes associated with muscle differentiation, metabolism, and contraction, while cell cycle-related genes were downregulated. In human skeletal muscle cells, Nr2f6 overexpression significantly increased the expression of myosin heavy chain genes (2-3-fold). Nr2f6 content in skeletal muscle decreased by 40% in lipid-overloaded myotubes and by 50% in mice fed a high-fat diet. Depletion of Nr2f6 increased myocyte lipid oxidative capacity by 75% and protected against lipid-induced cell death. This protection was associated with direct repression of uncoupling protein 3 (20%) and PGC-1α (30%) promoter activity following Nr2f6 overexpression. Nr2f6 overexpression in mice resulted in an atrophic and hypoplastic state, characterized by a significant reduction in muscle mass (15%) and myofiber content (18%), accompanied by an impairment (50%) in force production. These functional phenotypes were accompanied by the establishment of an immune response molecular signature and a decrease in genes involved in oxidative metabolism and muscle contractility. Additionally, Nr2f6 regulated core components of the cell division machinery, effectively decoupling muscle cell proliferation from differentiation.

**Conclusion:** In summary, our findings reveal a novel role for Nr2f6 as a molecular transducer that plays a crucial role in maintaining the balance between skeletal muscle contractile function and oxidative capacity. These findings have significant implications for the development of potential therapeutic strategies for metabolic diseases and myopathies.

## 1. Introduction

Muscle contraction is a highly coordinated process initiated by the transmission of the action potential from the afferent neurons to the muscle. Subsequent depolarization of the muscle fiber propagates along the sarcolemma, stimulating the release of calcium ions from the sarcoplasmic reticulum, and ultimately promoting myosin-actin cross-bridge cycling and force production. These processes require an accessory metabolic machinery to generate the energy needed to support contraction, and disruption in either muscle structure or metabolism leads to functional defects, such as in Duchenne syndrome [1], sarcopenia [2], and cachexia [3]. Dynamic crosstalk between energetic status, muscle development, mechanical stress, and transcriptional changes is crucial to maintaining muscle function. In this context, the nuclear receptor family of transcription factors (NRs) is of particular interest as they are potentially regulated by small molecules, such as metabolites and hormones [4]. Although the transcriptional landscape for metabolic-functional signaling has been extensively studied in pathological and physiological conditions, a broader role of some NRs has only recently been recognized, and thus, the role of many members remains elusive [5].

The NRs have a modular architecture displaying a ligand-binding domain (LBD) and a zinc-finger DNA binding domain (DBD) and can be further grouped in endogenous, orphan, or adopted NRs according to the presence of an endogenous ligand, the absence of a known ligand or if a new ligand for a given NR was just identified, respectively[6]. The classical mechanistic model proposes that a small molecule binds to the LBD, changing its conformation to one of higher affinity for a transcriptional co-regulator, such as PPARγ coactivator-1 alpha (PGC-1α), which in turn can mediate transcriptional modulation through the recruitment of histone modifier and the basal transcriptional apparatus [7]. The orphan nuclear receptor Nr2f6, also named Ear2, has been characterized in a broad range of tissues, such as adipose, thyroid, liver, brain, and the immune system, where it plays different and even antagonistic roles[8]. Nr2f6 can impair adipocyte differentiation, increase cancer cell proliferation, induce both resistance and susceptibility to antitumor drugs, and promote the development of fatty liver disease by upregulating CD36 [9]. So far, the most extensively defined role of Nr2f6 is in the immune system, in which it directly and strongly suppresses interleukins 17, 21, 2 and interferon γ transcription by interacting with the NFAT/AP-1 complex at the promoters of these genes [10], [11].

Curiously, Nr2f6 has been reported both as a transcriptional repressor and activator, but the context that defines its activity state is unknown. Recently, Nr2f6 was classified as a stripe transcription factor [12], i.e., it can bind to low-complexity motifs together with other transcription factors in a broad range of promoters, regulating chromatin accessibility. This suggests that Nr2f6’s function in different environments depends on its DNA occupancy and interaction with other transcription factors. As a result, it’s uncertain if the current understanding of Nr2f6’s role can be extended to other tissues like skeletal muscle. Therefore, we sought to characterize the molecular mechanisms and functional roles of Nr2f6 in skeletal muscle biology both *in vitro* and *in vivo*. Here, we report the functional and mechanistic characterization of Nr2f6’s role in the skeletal muscle. *In vitro* approaches identified Nr2f6 as a repressor of PGC-1α and the uncoupling protein 3 (UCP3) gene expression, consistent with the improvement in cells’ resilience to lipid toxicity and enhanced oxidative capacity in Nr2f6 loss-of-function models. Nr2f6 gain-of-function *in vivo* promoted an inflammatory transcriptional signature, and upregulation of core genes of cell cycle progression, culminating in muscle waste, change in fiber type, and impairment of force production.

## 2. Materials and Methods

The main reagents, tools, and models necessary for replicating the reported results are listed in Supplementary Table 1.

### 2.1. Cell culture

Human primary skeletal muscle cells were isolated from healthy female and male donors [13], age 55 ±5 years old, BMI 25.6 ±1.5 kg.m^-2^. Myoblasts were maintained in Growth Media (Dulbecco’s Modified Eagles Media (DMEM)/F12 High Glucose (Gibco, #31331093) supplemented with 10 mM HEPES (Gibco #15630-056), 16% Fetal calf serum (Sigma, #F7524), and antibiotics (Gibco #15240-062) and differentiated at the confluence with fusion media (74% DMEM High Glucose (Gibco, 31966-021), 20% 199 Medium (Gibco #31150-022), 20 mM HEPES, antibiotics, 0.03 µg/mL Zinc Sulfate (Sigma #Z4750), 1.4 mg/mL Vitamin B12 (Sigma #V6629), and 2% Fetal Calf Serum) supplemented with 100ug/mL Apotransferrin (Biotechne #3188-AT-001G) and 1.7 mM Insulin (Actrapid Penfill, Novo Nordisk #13509) before use. After 5 days of fusion, apotransferrin and insulin were removed from the media, and cells were incubated for 4 more days. Cells were cultivated in a humidified atmosphere containing 7.5% CO2 and regularly tested for mycoplasma. C2C12s, MEFs, and HEK cells were maintained in DMEM High Glucose (Gibco, 31966-021) supplemented with 4 mM L-glutamine, 10% fetal bovine serum, 1 mM sodium pyruvate, and antibiotics. Fetal bovine serum was substituted by 2% horse serum to induce myogenesis in C2C12 cells when 90-100% confluence was reached, and experiments were performed 5 days later in fully differentiated myotubes.

### 2.2. Primary mouse skeletal muscle cells

Mice’s primary skeletal muscle cells were isolated from wild-type C57Bl6/JUnib as described [14]. After euthanasia, hindlimb muscles were dissected and digested with collagenase II, trypsin, and DNAse I. Cells were sifted through a 70 µm cell strainer and plated in 0.1% Matrigel-coated plates. Myoblasts were maintained for 2 days in DMEM High Glucose supplemented with 2 mM L-glutamine, 10% fetal bovine serum, 10% horse serum, 1 mM sodium pyruvate, and antibiotics. Myogenesis was induced by removing fetal bovine serum from the media when total confluence was reached, and cells were cultivated for 5 more days to form fully differentiated myotubes. The experiments were approved by the Ethics Committee on Animal Use (CEUA/Unicamp #5626-1/2020).

### 2.3. Animals

All electroporation experiments were conducted following the guidelines of animal welfare and were approved by the Stockholm North Animal Ethical Committee (Stockholm, Sweden). Male C57Bl6/J mice were acquired from Jackson Labs and maintained at 12/12h light/dark cycle under controlled temperature and humidity, and *ad libitum* access to food (Specialized Research Diets, # 801722) and water. The use of animals for high-fat diet experiments was approved by the Ethics Committee on Animal Use (CEUA/Unicamp #5626-1/2020) and all the welfare guidelines of the National Council of Control of Animal Experimentation (CONCEA) were followed. Male C57Bl6/JUnib mice were kept under the same conditions described above. Mice were provided a high-fat diet (PragSolucoes #0015, 60% kcal from lipids) at 4 weeks of age for 16 weeks; littermates were fed a standard chow diet as a control.

### 2.4. Reactive oxygen species measurement

Cells were incubated with 5 nM MitoSOX (Invitrogen, #M36008) or 5 µM Dihydroethidium (DHE, Invitrogen, #D11347) for 30 min in DMEM without phenol red supplemented with 1 mM Sodium Pyruvate, 4 mM L-glutamine, and 25 mM Glucose and washed three times before reading in a plate reader 510/580 nm (ex./em.) or 520/610 nm (ex./em.) for MitoSOX and DHE, respectively. For normalization, samples were immediately fixed with 5% formaldehyde, stained with 0.05% Crystal Violet solution for 15 minutes, and thoroughly washed with water. The dye was resuspended in 10% acetic acid and absorbance read at 590 nm in a plate reader.

### 2.5. RT-qPCR

Total RNA was extracted from cells with TRIzol (Invitrogen #15596-018) following the manufacturer’s instruction and cDNA was synthesized with a High-Capacity Reverse Transcription kit (Applied Biosystems #4368814). cDNA was diluted to 10 ng/µL and 20 ng was used for qPCR reactions. NormFinder[15] was used to decide the best combination of internal controls among RPL39, PPIA, HPRT, 18S, ACTB, and GAPDH. In the *in vivo* electroporation experiments, gene expression was normalized using HPRT-PPIA geomean with TaqMan probes or HPRT-RPL39 geomean when SYBER green was used. For other experiments, gene expression was normalized with multiplexed HPRT when TaqMan probes were used or with RPL39 when SYBER was used. Relative gene expression was calculated by the ΔΔCT method[16] and is expressed as fold change over the indicated control. The primers and probes used are listed in Supplementary Table 2.

### 2.6. RNA-seq

Total RNA was extracted with TRIzol and the upper phase containing RNA was loaded into RNeasy columns (Qiagen, #74004) after the addition of isopropanol, following the manufacturer’s instructions. cDNA libraries were prepared with TruSeq Illumina Total RNA Stranded (Illumina) with Ribo-zero rRNA depletion (Illumina). Sequencing was outsourced to Macrogen Inc. (Seoul, South Korea) and performed in a HiSeq X (Illumina), producing an average of 50.8 Mreads, 95% above Q30. Sequence trimming and adapter removal were done with Trimmomatic[17] with the following modifications: HEADCROP = 10, MINLEN = 20, AVGQUAL = 20. Reminiscent reads were aligned to the mouse genome (Ensembl GrCm38) with RNA Star 2.7.2b and gene-level counts were calculated with featureCounts v1.6.4. Differential expression was performed with EdgeR with TMM normalization and p-value adjustment for multiple comparisons using Benjamini and Hochberg normalization with a 0.05 false discovery rate (FDR) cut-off. The Galaxy platform was used to process all data. Pathway enrichment analysis was done in g:Profiler with a 0.01 FDR cutoff. Interaction networks were generated by String.db and analyzed with CytoScape v3.8 using the EnrichmentMap plugin.

### 2.7. Fatty-acid treatment

Palmitate (Sigma, #P5585) in absolute ethanol and oleate (Sigma, #O7051) in water was conjugated with 1% fatty-acid-free bovine serum albumin (Sigma, #A7030) in cell media for 15 min at 55 °C to a 500 µM final concentration. Cells were treated with fresh solutions of palmitate or vehicle (1%BSA, 1% ethanol) for 20 hours.

### 2.8. Promoter transactivation assays

Luciferase reporter assays were performed in MEF cells transfected with a UCP3 reporter plasmid[18] (UCP3 EP1, Addgene #71743) or PGC-1α 2kb promoter (Handschin et al. 2003) (Addgene #8887), normalization plasmid coding for Renilla luciferase (pRL-SV40) and either control empty vector or Nr2f6 coding plasmid (Gift from Dr. Gottfried Baier, Medical University of Innsbruck, Austria) using Lipofectamine 3000. Luciferase activity was measured with DualGlo Luciferase Reporter Assay (Promega, #E2920). For knockdown assays, cells were transfected with siRNAs one day before the transfection of the reporter plasmids.

### 2.9. siRNA knockdown

C2C12 cells were transfected with 200 nM non-target siRNA (siScr, Qiagen) or siNr2f6 (Sigma) using Lipofectamine RNAiMax (Invitrogen) following manufacturer’s instructions concomitantly with the differentiation media switch. Experiments were performed on the third day of differentiation. Primary human skeletal muscle cells were double transfected, first concomitantly with the fusion media switch and later on the third day of differentiation with 5 nM siScr (Ambion) or siNr2f6 (Ambion). The experiments were performed on the seventh day of differentiation.

### 2.10. Stable cell lines

The Nr2f6-myc insert was subcloned from the Nr2f6-myc-flag plasmid into the pBABE-Puro vector using standard PCR with primers spanning the transcription start site and the myc tag. The viral particles for generating the overexpression HEK293T cells were transfected with pCMV-VSVG, pCL-Eco, and the pBABE-Nr2f6-myc or empty vector. For producing viral particles with the knockdown plasmid, HEK293T cells were transfected with packaging vectors pCMV-dR8.2 dvpr, pCMV-VSVG, and pLKO.1-shGFP or shNr2f6 (TRCN0000026147). Cell medium containing virions was collected, filtered at 0.45 µm, and stored at −80 oC until further use. Virus concentrations were titrated by the minimal dilution method, C2C12 cells were transduced with 1 MOI, and cells were selected with 2 µg/mL puromycin for 4 days. The clonal selection was performed in the knockdown cells and the clones were validated as indicated. The modified cells and their respective controls were cultivated synchronously under the same conditions.

### 2.11. Electroporation

Mice were kept under 2% isoflurane-induced anesthesia and the tibialis anterior muscles were injected with 30 µL of 1 mg/mL hyaluronidase (Sigma, #H3506). After 2 hours, the lateral and contralateral tibialis anterior were injected with 30 µg of either control empty vector pCMV6 or Nr2f6-myc-flag overexpression plasmid (Origene, #MR206083) and 220V/cm were applied in 8 pulses of 20/200 ms on/off (ECM 830 Electroporator, BTX). Terminal experiments were performed 9 days after electroporation with 13-week-old mice. For electroporation of FDB muscles, after anesthesia 10 µL of 1 mg/mL hyaluronidase were injected into the footpads and after 1 hour, 20 µg of the control or Nr2f6 coding plasmids. Mice rested for 15 minutes and then 75V/cm were applied in 20 pulses of 20/99 ms on/off with the aid of sterile gold acupuncture needles.

### 2.12. *Ex vivo* contraction

Mouse *flexor digitorum brevis* (FDB) muscles were electroporated as described and dissected 8 days later. To reduce variability dissections were performed by a single trained technician. With the muscles still attached to the tendons, contraction threads were tied at the most distal and proximal tendons, and the muscles were transferred to contraction chambers (Myograph system, DMT A/S) containing prewarmed and continuously oxygenated KHB buffer at 30 °C. The optimal muscle length was determined, and all subsequent measurements were performed at this length. For maximal force production mice, FDBs were stimulated at 10, 30, 50, 80, 100, and 120 Hz for 1 second and with 0.1 ms pulses. Muscles were left to rest for 5 minutes before starting the fatigue protocol as follows: 0.1 s train duration, 0.3 s train delay, and pulses of 0.1 ms at 50 Hz. The maximal force was evaluated again 5 min after the end of the fatigue protocol to check muscle integrity. Muscles were weighed and protein extraction was performed to normalize. The maximal force was calculated with the difference of the peak force at 120 Hz and the baseline and time to fatigue taken as the time necessary to reach 50% intensity of the first peak.

### 2.13. MHC Staining

Electroporated tibialis anterior muscles were embedded in O.C.T, immediately frozen in nitrogen-cooled isopentane, and stored at −80 °C until cryosectioning. Muscle slices were blocked (5% Goat serum, 2% BSA, 0,1% sodium azide in PBS) for 3 hours at room temperature and probed with primary antibodies overnight at 4 °C in a humidified chamber. The slides were washed 3 times with PBS and incubated with Alexa Fluor conjugated secondary antibodies (Invitrogen) for 2 hours at room temperature. Coverslips were mounted with ProLong antifade Diamond (Invitrogen) and whole sections were imaged with a fluorescent scanning microscope (AxioScan.Z1 Slide Scanner, Zeiss) at 20x magnification. Controls without primary antibodies were used to calibrate acquisition parameters and for image analysis. For fiber type quantitation, the images were randomized, and the analysis was performed blinded to the treatments, at least 2 consecutive cuts were quantified and averaged for each treatment-mice.

### 2.14. Oxygen consumption assays

Oxygen consumption rates (OCR) were measured in a Seahorse XF24 extracellular flux analyzer according to the manufacturer’s instructions. The following drugs were used in the assay: 1 μM oligomycin (Oligo), 2 μM carbonyl cyanate m-chlorophenyl hydrazone (CCCP), and 1 μM rotenone/antimycin (Rot./Ant). ATP-linked OCR was calculated by subtracting OCR post oligomycin addition from the OCR measured before. Reserve capacity was determined by subtracting basal from maximal OCR. Non-mitochondrial values were subtracted before all calculations. For fatty-acid oxidation assays, cell media was switched to low glucose 12 hours before the measurements, and cells were equilibrated in KHB supplemented with 1g/L glucose, 4 mM L-glutamine, and 1 mM sodium pyruvate for 1 hour. Immediately before the assay, BSA-conjugated palmitate was added to a final concentration of 200 µM, and the drugs were added in the same manner. During routine oxygen consumption assays, cells were maintained in phenol red-free DMEM, supplemented with 4.5g/L glucose, 4 mM L-glutamine, and 1 mM sodium pyruvate, without sodium bicarbonate.

### 2.15. Lactate measurement

Cells were grown in 96well plates and then incubated for 3 h with 50 µL Krebs-Henseleit Buffer (KHB) (1.2 mM Na2HPO4, 2 mM MgSO4, 4.7 mM KCl, 111 mM NaCl, pH 7.3) supplemented with 25 mM glucose, 1 mM pyruvate, and 4 mM Glutamine. Lactate production was enzymatically quantified as NADH fluorescence (360 nm/460 nm) by the reverse reaction of L-lactate dehydrogenase (Rabbit muscle, L25005KU, Sigma) in a reaction containing 20 µL cell media, 2 µg enzyme, 50 mM Tris, and 625 mM Hydrazine in PBS. Following the assay, the cells were fixed and stained with crystal violet for cell number normalization.

### 2.16. Western blot

Protein extracts from cells and tissues were obtained with RIPA Buffer (Thermo Scientific, # 89900). Briefly, cells were collected and homogenized in RIPA buffer with the aid of a sonicator, incubated on ice, and centrifuged at 16000 xg for 20 minutes to remove insoluble materials. Protein in the supernatant was determined using Bradford assay and 30 µg loaded into 4-10% or 4-12% gradient SDS-PAGE gels (Mini-PROTEAN TGX Precast Gels, BioRad). Proteins were then transferred to 0.45 µm PVDF membranes (Immobilon-P, Millipore) in a wet tank apparatus, probed with the indicated primary antibodies, and detected with ECL (ECL Select, Cytiva #RPN2235) in a ChemiDoc XR (BioRad). Images were analyzed with ImageLab software (BioRad) and protein band intensities were normalized by the Ponceau S intensity of the respective gel lane. Data is shown as fold change over control.

### 2.17. Microarray

RNA was extracted with TRIzol and subsequently column-purified using RNeasy Mini Kit (Qiagen). Sample integrity was assessed, and the library was prepared using an Affymetrix Whole Transcript (WT) Assay kit probed in a CGAS cartridge for Clariom S (mouse) following the manufacturer’s instructions. Total RNA quality was assessed by Agilent Technologies 2200 Tapestation and concentrations were measured by NanoDrop ND-1000 Spectrophotometer. Total RNA (150 ng) was used to generate amplified sense strand cDNA targets using GeneChip WT Plus Reagent Kit (ThermoFisher Scientific) followed by fragmentation and labeling. 2.3 µg of ss cDNA target was hybridized to Clariom S Mouse Arrays for 16 hours at 45 °C under rotation in Affymetrix Gene Chip Hybridization Oven 645 (ThermoFisher Scientific). Washing and staining were carried out on Affymetrix GeneChip Fluidics Station 450 (ThermoFisher Scientific), according to the manufacturer’s protocol. The fluorescent intensities were determined with Affymetrix GeneChip Scanner 3000 7G (ThermoFisher Scientific). Transcriptome Analysis Console (TAC) software (v4.0.3, ThermoFisher Scientific) was used for the analysis of microarray data. Signal values were log2-transformed, and quantile normalized using the Signal Space Transformation (SST-RMA) method. Since control and treated samples were obtained from the same animal, paired comparisons of gene expression levels between sample groups were performed using a moderated t-test as implemented in BioConductor package limma. Genes with FDR < 0.05 and fold change ≥ 2 were considered differentially expressed Gene ontology enrichment tests were performed with g:profiler excluding electronic annotations and using a significance threshold of 0.01.

### 2.18. Cell-death assays

Cell-death assays were performed as described [19] with slight modifications. Propidium iodide (Invitrogen #P3566) was added to a concentration of 5 µg/mL in cell culture media and incubated for 20 minutes. Hoechst 33342 (Invitrogen #H3570) was then added to a final concentration of 1 µg/mL and samples were incubated for another 10 minutes. Fluorescence was measured at 530/620 nm (ex./em.) and 350/460 (ex./em.) nm in a plate reader.

### 2.19. Cell doubling time

Cells (10^4^) were plated in four replicates in 12-well plates. Thereafter, cells were collected every 24 hours using trypsin and counted in a Neubauer chamber. The normalized data of three independent experiments were used to obtain the doubling-time regression curve with the initial number constraint.

### 2.20. ATP measurement

Cells were grown in opaque 96-well white plates and then processed according to the manufacturer’s instructions of the CellTiter-Glo Luminescent Cell Viability Assay kit (Promega # G9241). The standard curve of ATP was determined in parallel for absolute quantitation.

### 2.21. Bioinformatic analysis of public datasets

Nr2f6 ChIP-seq bigwig files from the ENCODE project available on GEO (Gene Expression Omnibus) accession codes GSM2797593 and GSM2534343 [20], [21] were used. Using the Galaxy platform, the anchoring position matrix was generated using the compute matrix command from the deeptools package, relative to human genome annotations extracted from the UCSC Genome Browser in bed format. The heatmap was obtained using the plotheatmap tool, also from the deeptools package. pathway enrichment was analyzed using the g.profiler program (https://biit.cs.ut.ee/gprofiler/gost), with a significance threshold of 0.01 using the g:SCS parameter, without considering electronic term annotations. The correlation in Supplementary Figure 1C was produced with fold changes of differentially expressed genes from RNA-seq (FDR <0.05) and fold change values (expression in myotube/expression in myoblast) from the C2C12 cell differentiation array (GSE4694) [22] considering a p-value cut-off of 0.01, according to GEO2R. UCP3 genome locus in Figure 3F was extracted from the UCSC genome browser with the ChIP-seq tracks of Myogenin (wgEncodeEM002136, wgEncodeEM002132), MyoD (wgEncodeEM002127, wgEncodeEM002129), H3K4me (wgEncodeEM001450), H3K27Ac (wgEncodeEM001450) and DNA hypersensitivity track (wgEncodeEM003399) over NCBI37/mm9 mouse genome assembly. Nr2f6 response elements searches (Fig. 3F and 6A) were performed using JASPAR motifs MA0677.1, MA0728.1, and MA1539.1 in RSAT matrix scanning with background estimated from the input sequence and 10^-5^ p-value cutoff.

### 2.22. Statistical analysis and quantification

GraphPad Prism v7.0 was used for plotting the data and for statistical analysis. Cell culture experiments were performed independently several times with at least 3 technical replicates in each independent experiment. Given the cells of individual donors of human skeletal muscle cells are kept independently and each represents a single person, comparisons were made between the control and treated samples within the same donor using paired (ratio) comparison Student’s t-test was used. The same principle was used for electroporation experiments, in which control and treated samples come from the same mice, otherwise, unpaired comparisons were performed and a 0.05 p-value significance cutoff was used. Before all comparisons, data normality was confirmed by the Shapiro-Wilk test. Details for microarray and RNA-seq statistics are described in their respective methods section and further statistical details are provided in figure labels.

## 3. Results

### 3.1. Nr2f6 regulates the transcriptional landscape of myogenesis and metabolism in skeletal muscle cells

Genetic manipulations of Nr2f6 at the whole-body level and *in vitro* have been conducted[9], [23]–[25], but the role of this nuclear receptor in muscle models is underexplored. We used siRNA-mediated depletion of Nr2f6 in C2C12 myocytes (Fig. S1A) to verify the outcomes on the transcriptomic landscape. The 1849 differentially regulated genes, 920 upregulated and 939 downregulated, could be grouped into five main classes, with increased expression of genes related to muscle differentiation, contraction, and metabolism and decreased expression of genes with roles in cell cycle and DNA packaging (Fig. 1A, B, E). In fact, among the 20 most significant differentially expressed genes, eleven are linked to muscle contraction (*RYR1, TTN, MYH3, MYH2, ACNT2, ATP2A1, MYL1, TNNC2, MYOM3, CACNA1S, LRP4*) and other three compose the cytoskeleton (NEB, MACF1, XIRP1), all upregulated in the Nr2f6 knockdown (Fig. S1B). Accordingly, a panel of canonical markers of muscle differentiation containing muscle regulatory factors (MRFs) and myosin isoforms (Fig. 1C) shows a broad upregulation of these indicators. Indeed, data generated in our transcriptomic analysis correlated directly with the publicly available C2C12 myogenesis dataset (Fig. S1C), indicating an association between Nr2f6 levels and myogenic potential. Consistently, the proteins coded by the upregulated genes belonged mostly to the sarcomere, contractile fiber, and cytoplasm location ontologies. Myogenic differentiation demands the withdrawal of the cell cycle and both processes are regulated in a concerted manner [26], [27].

**Figure 1.**
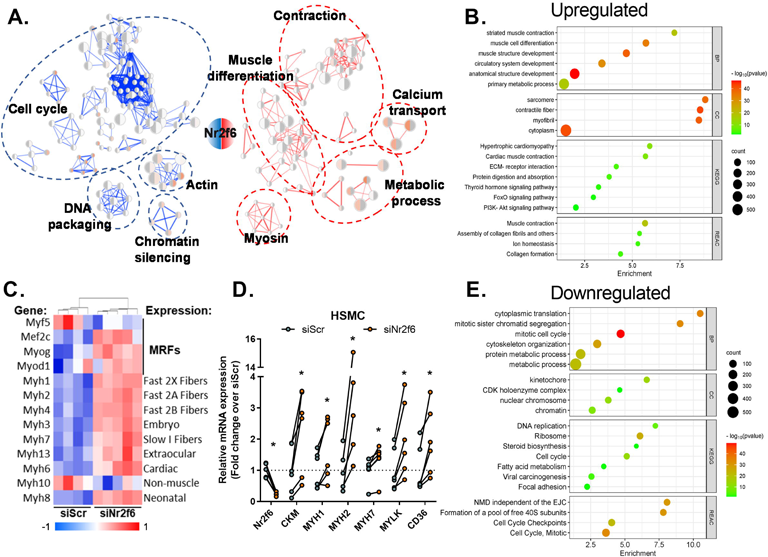
Nr2f6 knockdown derepresses the expression of genes involved in metabolism and myogenesis. (A) Network of ontology terms enriched in the differentially expressed genes in the transcriptomics of transient Nr2f6 knockdown in C2C12 cells. Groups of similar terms were manually curated and encircled as indicated. Upregulated elements in red and downregulated in blue (N=4-5). (B, E) Gene ontology enrichment of downregulated and upregulated genes. (C) Panel of myogenic differentiation markers differentially regulated by Nr2f6 knockdown with Myogenic Regulatory Factors (MRFs) and myosin isoforms and their respective fiber expression patterns [90], [91]. (D) Gene expression measured by RT-qPCR of markers of myogenic differentiation in primary human skeletal muscle myotubes (HSMC) transfected with control non-target RNAi (siScr) or siNr2f6 (N=5-6). Circles represent individual donors. * Indicates p < 0.05 using paired two-tailed Student’s t-test.

Accordingly, the downregulated genes were related to distinct phases of cell division, with the enrichment of proteins related to DNA replication, packaging, and chromosome segregation. Of note, essential components of the cell cycle progression such as *CDC25B/C* phosphatases and *CDK1/4* kinase promote quiescence and halt cell cycle progression in muscle progenitors when down-regulated[28]–[30]. Considering the differences between mouse and human myogenic cell’s transcriptional landscape[31] we verified whether the result of Nr2f6 depletion in differentiation markers would be translatable to human primary human skeletal muscle cells. We found that expression of *MYH1/2/7*, muscle creatine kinase, and myosin light chain kinase 1 were upregulated in human myotubes (Fig. 1D), indicating a conserved role for Nr2f6 as a repressor of myogenesis. Since both an increase in cellular oxidative capacity and the activation of the PI3K pathway are required for the completion of myogenesis[32], [33], we verified whether the genes of major pathways of regulation of glycolysis and fatty acid oxidation were affected and found that several energy sensors such as *AKT2*, *PRKAG3* subunit of AMP-activated protein kinase (AMPK) and the mTOR complex were upregulated by Nr2f6 loss-of-function (Fig. S1D). Altogether, the changes in the myocyte’s global transcriptome indicate that Nr2f6 represses key gene networks associated with myoblast differentiation and metabolism.

### 3.2. Nr2f6 depletion improves oxidative metabolism and enhances lipid handling capacity

Given the enrichment of genes of oxidative metabolism, we first sought to verify the functional effects of Nr2f6 knockdown on metabolism. Oxygen consumption assays (Fig. 2A, B), using palmitate as the major substrate for energy production, provided evidence that Nr2f6 knockdown in C2C12 myocytes promoted an increase in maximal respiration and spare capacity after the addition of the uncoupler carbonyl cyanide m-chlorophenyl hydrazone (CCCP). Although there was no difference between control and knockdown respiratory parameters in high-glucose media (Fig. S2A, B), the extracellular acidification rates were reduced, without a reduction of total ATP pool, which was confirmed by lower lactate concentration in knockdown cells (Fig. 2C, S2C, D). Together, these results suggest that Nr2f6 knockdown increases oxidative metabolism. To assess the effects of sustained Nr2f6 depletion on muscle cell phenotype, we established a stable C2C12 cell lineage with a 70% reduction in Nr2f6 expression through retroviral knockdown (Fig. 2E). In line with the metabolic alterations, Nr2f6 knockdown increased the expression of the glucose transporter *GLUT4*, the anaplerotic enzyme pyruvate carboxylase (PC), and fatty-acid transporters (Fig. 2E). Our analysis of ENCODE chromatin immunoprecipitation-sequencing (ChIP-seq) data for Nr2f6 in K562 and HepG2 cells indicates an increase in mitochondrial genes related to lipid metabolism (Fig. S2G, H). We also identified significant enrichment of kinases related to the insulin signaling pathway when analyzing genes affected by Nr2f6 knockdown in the RNA-seq data and genes with Nr2f6 binding peaks within the promoter region in ENCODE with ChIP-seq data (Fig. S1H). Considering the increase in the efficiency of the Nr2f6-silenced myocytes to oxidize lipids, we hypothesized that depletion of Nr2f6 would protect against lipid overload in skeletal muscle. Stable Nr2f6 knockdown in myotubes attenuated palmitate-induced cell death (Fig 2F) and prevented the increase in mitochondrial and cytosolic superoxide production induced by palmitate overload(Fig. 2D). Next, we verified whether Nr2f6 is modulated by palmitate treatment *in vitro* and by a high-fat diet *in vivo* in rodents, as models of increased lipid oxidation and supply[34]. Our findings demonstrate that Nr2f6 expression is consistently reduced under these conditions, indicating a role as an energy stress response gene that facilitates metabolic adaptations to lipid oxidation (Fig. 2G-I), S2E, F). Collectively, the data provide evidence that Nr2f6 inhibition protects against lipid overload by increasing lipid handling capacity in skeletal muscle in stress conditions, possibly by a concerted upregulating of mitochondrial and cytosolic lipid transporters, mitochondrial proteins, and TCA cycle anaplerotic genes, thereby increasing oxygen consumption.

**Figure 2.**
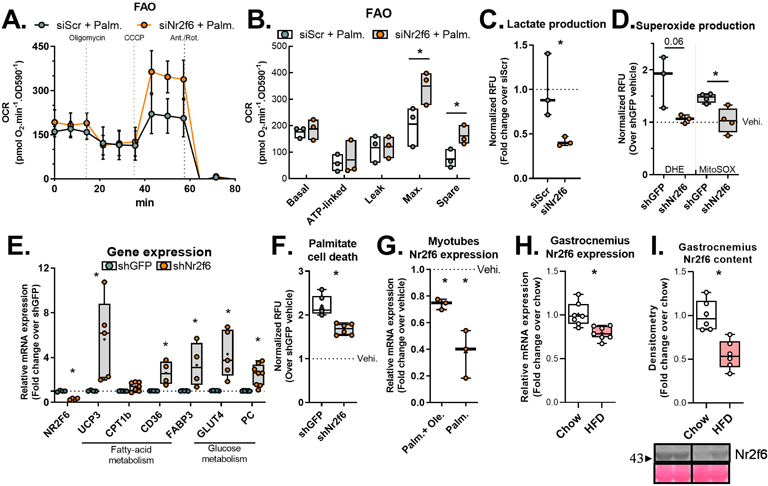
Nr2f6 depletion increases fatty acid oxidation and protects cells against lipid-induced stress. (A) Fatty-acid-dependent oxygen consumption (FAO) assay in control siScr and siNr2f6 C2C12 myocytes using palmitate (palm.) as substrate. Data displayed as mean ±SD. (B) Calculated respiratory parameters of the FAO assay are displayed as a line on the mean and minimum to max bars (N=3). * Indicates p < 0.05 using unpaired two-tailed Student’s t-test. (C) Lactate measurement in cell culture media of C2C12 myocytes transfected with control siScr and siNr2f6. (N=3). (D) Mitochondrial and total superoxide production following palmitate treatment in shGFP and shNr2f6 stable C2C12 cells (N=3-4). (E) Relative gene expression using RT-qPCR in stable Nr2f6 knockdown (shNr2f6) C2C12 cells and control shGFP stable cells (N=4-8). (F) Cell death as measured by propidium iodide in control (shGFP) and shNr2f6 myocytes following treatment with 500 µM palmitate for 20 hours (N=3). (G) Relative Nr2f6 mRNA expression in C2C12 myotubes treated with 500 µM and 500 µM oleate (Ole.), 500 µM palmitate (Palm.), or vehicle for 20 hours (N=3). (H, I) Relative Nr2f6 mRNA expression (N=7) and protein content (N=6) in the gastrocnemius of mice fed with a control chow diet or high-fat diet for 16 weeks. Below: representative western blot image. Boxplot with whiskers spanning minimum to maximal and box edges 25th-75th percentile, the line at the median and + at the mean. * Indicates p < 0.05 using unpaired two-tailed Student’s t-test.

### 3.3. Transrepression of PGC-1α and UCP3 transcription by Nr2f6

We recently provided evidence that the transcriptional regulation of *UCP3* by the peroxisome proliferator-activated receptor γ co-receptor 1-α (PGC-1α) is essential for the maintenance of myotube viability during lipid overload, by sustaining oxidative capacity in both acute and chronic treatment [35]. Considering a similar phenotype by Nr2f6 knockdown in myotubes, we investigated whether Nr2f6 regulates the same pathway. We found that Nr2f6 overexpression (Fig. 3SA, B) reduced both *UCP3* and *PGC-1α* mRNA in myotubes, which also translated in a decrease of PGC-1α protein content and expression of its mitochondrial electron transfer chain (ETC) target genes (Fig. 3A, B). Conversely, Nr2f6 knockdown in C2C12 myotubes increased PGC-1α downstream ETC targets (Fig. 3C). Reinforcing an upstream action of Nr2f6 over PGC-1a transcription, luciferase reporter assays showed an inhibition in promoter transactivation suggesting direct repression (Fig. 3D). Subsequent investigations of the regulation of *UCP3* transcription by Nr2f6 using 7kbp UCP3 promoter reporter plasmid demonstrated a decreased luciferase signal in Nr2f6 overexpression and an increased activity following Nr2f6 knockdown (Fig. 3E). Scans of UCP3 promoter with Nr2f6 motif found an Nr2f6 response element downstream of the transcription initiation site, which coincided with open chromatin region and peaks of known the UCP3 transcription factors, Myod1 and Myogenin (Fig. 3F), further supporting the notion that Nr2f6 directly repressed *UCP3* expression. Importantly, the repressive effects of Nr2f6 on *UCP3* and *PGC-1α* expression were conserved in primary myotubes of both humans (Fig. 3G) and mice (Fig. S3C). UCP3 transcription is regulated by peroxisome proliferator-activated receptors (PPARs) and estrogen-related receptors (ERRs) in skeletal muscle[36], [37], however, Nr2f6 silencing did not change the transactivation of responsive elements (Fig. S3D). Collectively, our results indicate that Nr2f6 is a bona fide transcription regulator of UCP3 and PGC-1α. Given that UCP3 is a PGC-1α target, these results indicate that Nr2f6 represses UCP3 expression indirectly through Pgc-1α downregulation and by modulation of UCP3 promoter activity.

**Figure 3.**
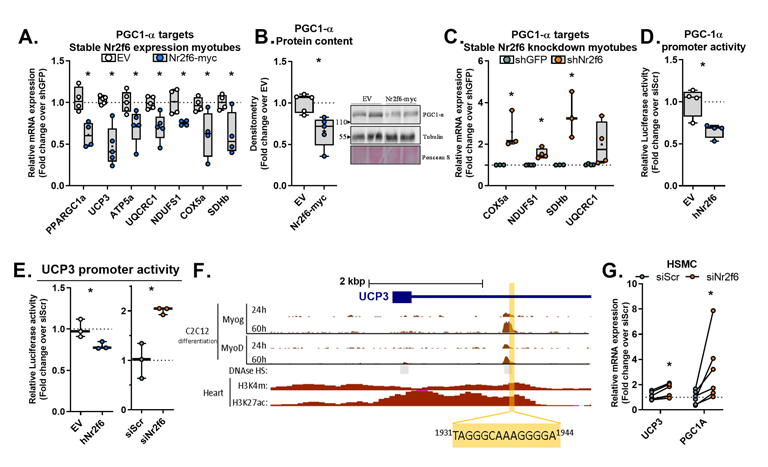
Nr2f6 inhibits PGC1-*α* and UCP3 gene expression. (A) Relative gene expression using RT-qPCR in stable Nr2f6-myc overexpression myotubes (N=4-5). (B) Densitometry and representative images of PGC-1α western blot in stable Nr2f6-myc overexpression myotubes (N=5). (C) Relative gene expression by RT-qPCR in stable Nr2f6 knockdown myotubes (N=3-5). (D) PGC-1α 2kbp luciferase reporter assay in HEK293 cells overexpressing HA-tagged Nr2f6 (Nr2f6-HA) or control empty vector (EV). (E) Luciferase activity of UCP3 promoter transactivation assay in cells transfected with Nr2f6-HA or siNr2f6 (N=3). (F) Mouse UCP3 genomic locus retrieved from UCSC Genome Browser with the Nr2f6 response element highlighted. Top tracks: ChIP-seq of Myogenin and MyoD at 24h and 60h of differentiation. Middle track: DNAse hypersensitivity assay, with open sensitive regions in grey. Bottom tracks: histone marks ChIP-seq. (G) Relative gene expression by RT-qPCR in human primary skeletal myotubes (HSMC) transfected with siNr2f6 or siScr (N=6). Boxplot with whiskers spanning minimum to maximal and box edges 25th-75th percentile, the line at the median and + at the mean. * Indicates p < 0.05 using paired two-tailed Student’s t-test for the experiments with human cells and unpaired for the other comparisons.

### 3.4. Nr2f6 promotes cell proliferation and represses genes implicated in muscle contraction and oxidative metabolism

We next explored the effects of Nr2f6 overexpression *in vivo* by electroporation in the *tibialis anterior* muscle of mice, using the contralateral muscle as control, and studied the global transcriptomic changes by microarray. There were 3796 genes within the criteria for differential expression (FDR <0.05, fold change >2), among which, 1915 were downregulated and 1781 were upregulated, with Nr2f6 overexpression having a major effect on the hierarchical clustering (Fig. S4A). Consistent with reports that highlight Nr2f6 as a gatekeeper of the immune system[24], gene ontology analysis of the upregulated genes shows enrichment of biological processes and pathways related to the immune system (Fig. 4B). Further validation by RT-qPCR indicated that markers of lymphocyte activation *CD44*, the marker for macrophage/monocyte activation *CD68*, and the macrophage marker *F4-80* were upregulated in Nr2f6 expressing muscle (Fig. 4E). Accordingly, *TGFb*, a potent inhibitor of hematopoietic cell activation, and the marker for endothelial and non-differentiated hematopoietic cells were downregulated. These results suggest that Nr2f6 might activate resident cells of the immune system and/or promote the invasion of circulating cells. Nr2f6 overexpression also increased expression of Myogenin and, to a lesser extent, *MYOD*, however the downstream targets genes myosin heavy chains 1 and 2 decreased (Fig. 4D). Downregulated genes were enriched in energetic metabolism, mitochondria, and muscle contraction terms (Fig. 4C), consistent with the functional phenotypes described *in vitro*. Importantly, the hereby proposed Nr2f6 targets, namely *UCP3* and *PGC-1α*, were also repressed by Nr2f6 *in vivo* (Fig. 4F). The lipid transporters CD36 and CPT1B, and subunits of the respiratory chain complexes, upregulated by Nr2f6 knockdown in vitro, were also downregulated by Nr2f6 overexpression. Additionally, the expression of reactive oxygen species scavengers *SOD1*, *SOD2*, and catalase genes was decreased (Fig. 4F). Collectively, these findings further confirm our functional results *in vitro* and indicate mitochondrial activity was impaired by Nr2f6 overexpression.

**Figure 4.**
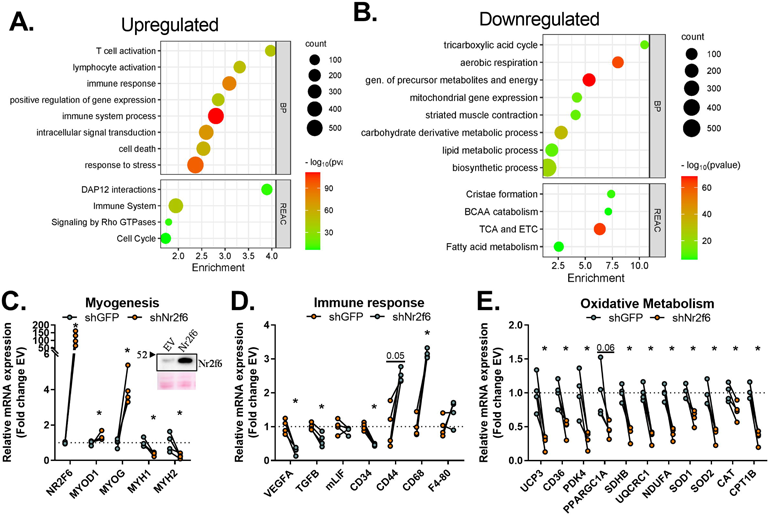
The transcriptional landscape of Nr2f6 overexpression exhibits a decrease in metabolism and an increase in inflammatory markers. (A, B) Gene ontology enrichment of downregulated and upregulated genes. (D, E, F) Validation of selected markers modulated in the microarray by RT-qPCR. N=4. Insert on D: representative western blot for validation of Nr2f6 overexpression in the *tibialis anterior* samples under control (EV) and electroporated(Nr2f6-myc) muscles. Circles represent individual samples. * Indicates p < 0.05 using paired two-tailed Student’s t-test. The numbers above some bars indicate the p-value.

### 3.5. Nr2f6 overexpression intensifies muscle waste and impairs force production

Since Nr2f6 overexpression negatively affects the mRNA expression of genes involved in muscle contraction and development, we considered whether the Nr2f6 gain-of-function would impair muscle morphology and function by performing immunostaining for myosin heavy chain (MHC) isoforms and *ex vivo* contraction experiments. Strikingly, Nr2f6 overexpressing *tibialis anterior* (TA) weighted less and were visually paler compared with control muscle (Fig. 5A). Consistent with these observations, Nr2f6 overexpression reduced (20%) the total number of fibers (Fig. 5B, C), which together with the increase in cell death-related genes (Fig. S4B), and the increase in the atrogenes cathepsin and calpain 2[38] (Fig. S4B), characterizes a state of atrophy and hypoplasia. Stratification by fiber type showed that this reduction is particularly due to the decrease (25%) in type IIB fibers, and although there was a tendency to decrease IIX fiber number (Fig. 5D), there was no statistical significance in these comparisons or the number of IIA fibers. To investigate whether these molecular disturbances result in functional alterations, we overexpressed Nr2f6 in the *flexor digitorum brevis* (FDB) and performed *ex vivo* contractions. Considering the length constraints for ex vivo contraction, the FDB muscle provided the ideal model for these experiments, due to its small size, ease of access for electroporation, and similarity with TA fiber composition, consisting mostly of type II fibers[39]. Consistently, mass-corrected maximal force production was considerably reduced (60%) in Nr2f6 overexpressing muscles (Fig. 5E). No effects were observed in the time to fatigue between control empty vector and Nr2f6 overexpression (Fig. 4SC). The *ex vivo* contraction assay surpasses the neuromuscular system by direct electric stimulation, disregarding effects on action potential transmission. In this setup, fatigability is mostly induced by detriments in calcium homeostasis, such as reduced Ca^2+^ sensitivity, sarcoplasmic Ca^2+^ reuptake, and release[40], therefore we cannot exclude the possibility that the time to fatigue is also affected *in vivo* in Nr2f6 gain-of-function models. Altogether, these findings strongly suggest that Nr2f6 induces muscle loss, worsened by an overactivation of the immune system state and an imbalance between satellite cell proliferation and differentiation.

**Figure 5.**
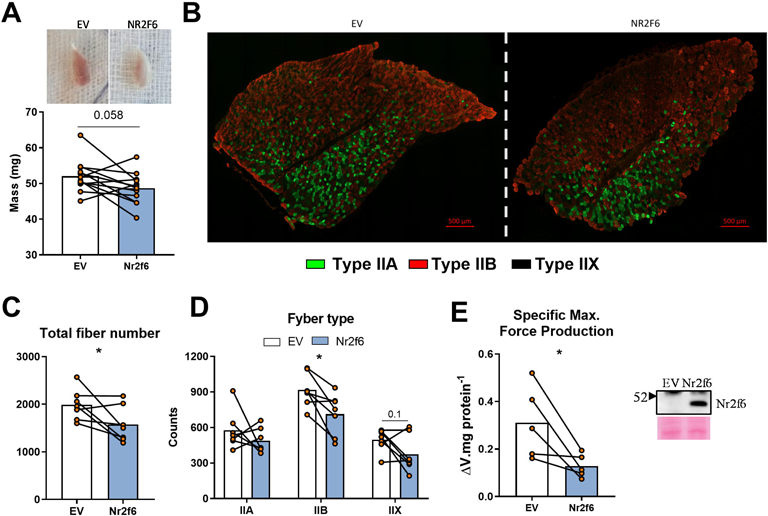
Overexpression of Nr2f6 induces muscle atrophy and impairs muscle force production. (A) Weight of *tibialis anterior* muscles (TA) electroporated with empty vector (EV) or Nr2f6-myc coding plasmid. Top: representative photo of electroporated muscles. N=12. (B) Representative images of myosin heavy chain staining in the electroporated TAs for fiber type determination. In green, MHC IIA; in red, MHC IIB; unstained fibers as IIX. No substantial number of MHCI fibers were stained, therefore the corresponding channel was omitted. N=7. (C, D) Total and type-segmented fiber counts. N=7. (E) *Ex vivo* contraction maximal force production in FDB muscles electroporated with control empty vector (EV) or Nr2f6-coding plasmid. N=5. Data are displayed as individual animals and bars at the mean. * Indicates p < 0.05 using ratio paired two-tailed Student’s t-test.

**Figure 6.**
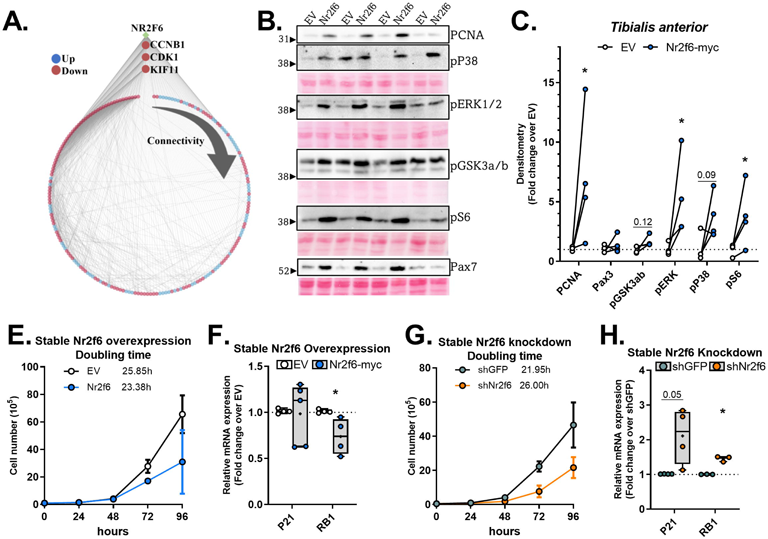
Nr2f6 increases myoblast proliferation rates. (A) Interaction network of genes consistently regulated by Nr2f6 overexpression and knockdown and with detected Nr2f6 binding motif at the promoter region. In blue: genes downregulated; in red: genes upregulated. The number of connections of each gene increases clockwise. (B, C) Representative images of the western blot of control (EV) or Nr2f6-myc electroporated *tibialis anterior* muscles and densitometric quantitation of the indicated protein bands. N=4. (D, F) Proliferation curves of stable C2C12 cell lines and the calculated doubling time. (E, G) RT-qPCR of cell cycle arrest markers in Nr2f6 knockdown and overexpression stable cell lines, respectively. N=4-6. Boxplot with whiskers spanning minimum to maximal and box edges 25th-75th percentile, the line at the median and + at the mean. * Indicates p < 0.05 using paired two-tailed Student’s t-test for mice experiments or unpaired for the other comparisons. The numbers above some bars indicate the p-value.

### 3.6. Myoblast proliferation rates are governed by Nr2f6

Next, we investigated whether Nr2f6 regulates cell cycle genes and myoblast proliferation. Thus, we compared differentially expressed genes identified in the microarray of the Nr2f6 overexpression in TA muscle with the RNA-seq transcriptomics from C2C12 myocytes after transient Nr2f6 knockdown. We found 706 genes were differentially expressed in both experiments, whereby 446 genes were modulated in opposite directions indicating a direct regulation by Nr2f6 or a conserved effect of Nr2f6 modulation in both models (Fig. S5). We further scanned the promoter regions (± 3kbp of the transcription start site) with the three Nr2f6 binding motifs available on the JASPAR database. We found matches in 206 unique genes, whereby 73 were upregulated and 133 downregulated by Nr2f6 overexpression. The interaction network of these high-confidence targets (Fig. 6A) revealed that the most connected genes are upregulated by Nr2f6 overexpression and integrate cell cycle pathways, which emphasizes the role of Nr2f6 as a promoter of cell division, and further reinforces the dysplastic phenotype observed in the gain-of-function experiments *in vivo*. Investigation of markers of myogenesis cell proliferation and stemness in Nr2f6 overexpressing TAs (Fig. 6B, C) indicated that the content of the proliferating cell nuclear antigen (PCNA), a fundamental marker for cell proliferation[41], was increased by Nr2f6 overexpression. Together with the increase of the muscle-specific satellite cell marker Pax7, and the activation of signaling cascades of the stemness marker GSK3a/b [42], [43], and proliferation markers ERK and p38 MAP kinases [44], [45], and S6[46], [47], these results point to an increase in myogenic progenitors and the infiltration of other cell types, such as cells of the immune system. Nr2f6 overexpression and knockdown can promote or inhibit cell proliferation in cancer cells, respectively [48]–[50]. Doubling time experiments confirm that this effect is also conserved in C2C12 myoblasts, with an increase (4 h) in the average doubling time by Nr2f6 depletion and a decrease (3.5 h) by Nr2f6 overexpression (Fig. 6D, F). Moreover, RT-qPCR validation of major markers of cell cycle progression inhibitors *RB1* and *P21* show an increase in both genes by Nr2f6 knockdown and repression of *RB1* by Nr2f6 overexpression (Fig. 6E, G). Collectively, these results demonstrate that Nr2f6 works as a major repressor of cell cycle progression in the skeletal muscle, directly modulating transcriptional networks implicated in cell division.

## 4. Discussion

Numerous nuclear receptors are necessary for the maintenance of muscle mass [51], [52]. For example, whole-body knockout of Nr1d1 (Rev-ERBα, Ear-1) leads to an increase in atrophic genes, a decrease in muscle mass, and a relative increase in low-diameter fibers[53]. More broadly, muscle-specific knockout of the nuclear receptor co-repressor 1 (NCoR1) leads to skeletal muscle hypertrophy and increased oxidative metabolism [54]. Here, using several skeletal muscle models we found evidence that Nr2f6 overexpression disrupts myogenesis *in vivo* and *in vitro,* and activates myoblast proliferation. Remarkably, Nr2f2 (COUP-TFII), an Nr2f6 interactor, is among the few nuclear receptors known to promote muscle wasting [55], [56]. Nonetheless, Nr2f2 expression in myogenic progenitors impairs muscle differentiation in mice by directly repressing genes related to both myoblast fusion and proliferation [57], [58], implying that these nuclear receptors regulate distinct phases of muscle differentiation. Future studies should address the redundancy of these NRs in muscle function.

Most of the transgenic and knockdown models provide evidence to suggest that nuclear receptors are involved in a general activation of oxidative metabolism[51]. For example, the muscle-specific Nr4a3 transgenic mouse displays a marked increase in mitochondrial density and fast-to-slow fiber switch [59]. This general rule is reinforced by the fact that mice lacking nuclear receptor coactivators, such as PGC-1α and MED1 [60], or overexpressing the co-repressor RIP140 [61], have decreased mitochondrial density and fewer oxidative fibers. Conversely, here we show that Nr2f6 is an exception to this model in skeletal muscle since it downregulates UCP3 and PGC1-α expression in human and mouse cells by transrepression of promoter activity. This reduces the myocyte’s capacity to oxidize fatty acids and increases reactive oxygen species production, leading to a higher susceptibility to lipotoxicity. Transgenic mouse models overexpressing UCP3 in muscle are consistently reported to have improved glucose homeostasis under chow and high-fat diet (HFD) conditions, as well as resistance to obesity-induced diabetes [60]–[62]. Moreover, increased levels of circulating lipids increase UCP3 expression [65], [66]. Here, we demonstrate that direct fatty-acid exposure in C2C12 myotubes or conditions of increased β-oxidation *in vivo* reduces Nr2f6 mRNA expression and protein content, lifting the repression on the UCP3 promoter, further reinforcing the finding that Nr2f6 mediates the positive effects of UCP3 on lipid handling under physiological conditions and that Nr2f6 downregulation is part of an adaptative response to lipid exposure.

Sarcopenia is the age-related loss of muscle mass and function, with the reduction of the number and size of myofibers, a switch from type II to I fibers[67], and an underlying mitochondrial dysfunction [68]. This henotype is closely reproduced by Nr2f6 overexpression in skeletal muscle. In contrast to most members of the NR family, Nr2f6 overexpression not only induces a sarcopenic-like phenotype, with atrophy, hypoplasia, and inflammation but also reduces muscle strength and affects energy metabolism. Interestingly, Nr2f6 is upregulated 2-fold in skeletal muscle hereditary spastic paraplegia [69], a disease characterized by progressive lower limb muscle weakness, sometimes accompanied by mitochondrial dysfunction and morphological fiber defects [47], [48]. Nr2f6 knockdown in myotubes increases myosin heavy chain expression and improves mitochondrial function, suggesting that inhibition of Nr2f6 might be efficacious in the treatment of sarcopenia or other myopathies. These findings are in line with recent bioinformatic analysis indicating Nr2f6 as a putative regulator of muscle energetic balance and development[72].

Additional experiments to assess the effects of Nr2f6 ablation in clinically relevant models are necessary to address this matter. Conversely, Nr2f6 agonists have been proposed as a treatment for colitis [73], however, in light of our results indicating a considerable muscle waste in Nr2f6 gain-of-function, the use of such agonists should be further evaluated for specific groups of patients suffering from myopathies and cachexia. Considering the known effects of Nr2f6 in the immune response, the atrophic phenotype described here could be partially underlaid by the action of Nr2f6 in immune cells. Nonetheless, the antagonistic transcriptional and functional changes induced by Nr2f6 overexpression *in vivo* and its knockdown *in vitro* in various experimental models strongly indicate a direct action of Nr2f6 in the myofibers as the major driver of the functional changes. A conceivable model elucidating the mechanism by which Nr2f6 overexpression culminates in a reduction of muscle force production (Fig. S4D), entails the downregulation of genes engaged in various facets of muscle contraction, such as muscle structure, calcium cycling, and action potential. Notably, a few of these genes constitute high-confidence targets. The ryanodine receptor 1 (*RYR1*) is a major component of the calcium release complex, which permits calcium efflux from the sarcoplasmic reticulum into the cytosol. As such, *in vivo* knockdown, and mutations of *RYR1* can cause severe myopathies[74]. Other putative targets including myosin light chain kinase 4 (*MYLK4*) and myomesin 1 (*MYOM1*) are downstream of the androgen receptor (AR), which mediates the effects on muscle force production [75]. Interestingly, we found that the spermine oxidase gene (*SMOX*), another important target of the AR in muscle [76], [77] is downregulated by Nr2f6 overexpression *in vivo*, upregulated by Nr2f6 silencing in C2C12 myocytes, and contains Nr2f6 binding motifs at the promoter region. The direct link between Nr2f6 and the spermine synthesis pathway and a possible interaction with the AR warrant further studies.

Targeted studies provided evidence that Nr2f6 activates gene expression by tethering to the promoters of *circRHOT1*[78], *DDA1* [79], and *CD36* [9], and represses the expression of numerous others such as *IL17*, *IL21*, Renin, and Oxytocin [48], [49], [80], [81]. More broadly, our transcriptomics experiments display an equilibrated number of genes up-and downregulated, further implying that Nr2f6 is a dual-function transcription factor. Interestingly, Nr2f6 is a target of MiR-142-3p[82], raising the possibility that miRNAs, besides protein partners and post-translational modifications[48], might aid in the regulation of Nr2f6 activity. Further studies should help to elucidate the mechanism by which Nr2f6 acts as a repressor or activator of gene expression. The case of CD36 illustrates context-dependent regulation given that this gene is activated in the liver [9], but repressed in skeletal muscle by Nr2f6 in mice and humans. This finding suggests that Nr2f6 may be bound to DNA but kept in a repressive state by post-translational modifications or interaction partners, such as RAR Related Orphan Receptor γ (RORγ), another regulator of *CD36* transcription in muscle[83] and liver[84]. In Th17 lymphocytes, Nr2f6 can compete with RORγ for binding at the *IL17* promoter, maintaining the repressive state. Our results support the proposition that this relationship could be conserved in the skeletal muscle.

In ovarian cancer cells, Nr2f6 tethers the histone acetylase P300 to the Notch3 promoter, which activates its transcription by increasing histone H3K9 and K27 acetylation and increases cell proliferation (4). We found that Nr2f6 also modulates myoblast proliferation *in vitro* and increases the expression of proliferation markers such as *PCNA* and *KI67 in vivo*. Additionally, key regulators of the cell cycle, such as cyclin B1 (Ccnb1) and Cdk1, are placed among high-confidence direct targets of Nr2f6. Ccnb1 interaction with Cdk1 is necessary for its kinase activity[85] and progression through the G2 mitotic phase. Ccnb1 expression is increased in several cancer types and ectopic expression increases cell proliferation rates [86], [87]. The simultaneous effect of Nr2f6 modulation on cell cycle and differentiation markers might also be sustained indirectly by the expression of the retinoblastoma protein, which is responsible for cell cycle arrest through the inhibition of the E2F family of transcription factors. In a feedback loop, E2F transcription factors antagonize MyoD1, which induces Rb expression, thereby linking processes of proliferation and differentiation[88], [89]. While Nr2f6 increases the expression of Myogenin in the *tibialis anterior* muscle, myosin heavy chains and other indicators of terminally differentiated myofibers are sharply reduced. These findings raise the possibility that Nr2f6 overexpressing myoblasts proliferate but fail to assemble into robust myofibrils due to a dysregulated modulation of the MRFs during myogenesis.

The evidence gathered here points to a global regulation of pivotal aspects of skeletal muscle biology by Nr2f6, through a concerted control of multiple gene networks. Nr2f6 modulation alone can determine myoblast proliferation rates, consolidating its role as a general regulator of cell cycle progression in different lineages, in skeletal muscle acting as a fulcrum between myogenesis and cell division. Conversely, the metabolic outcomes of Nr2f6 control of transcription are tissue-specific, and our findings show that lifting the direct repression of major players of mitochondrial activity leads to an improvement in muscle oxidative capacity. Our findings establish Nr2f6 as a novel regulator of muscle contraction and metabolism. This discovery suggests a potential avenue for its application as a prospective therapeutic strategy in addressing conditions characterized by muscle wasting disorders and metabolic diseases.

## Author Contributions

Conceptualization, D.S.P.S.F.G., N.M.F.B, A.G.O., D.R.R., M.J., J.A.B.S., T.R.A., A.S.V., A.K., J.R.Z., L.R.S.; Methodology, D.S.P.S.F.G., N.M.F.B, D.R.R., M.J.; Validation, D.S.P.S.F.G., N.M.F.B, L.R.S.; Formal Analysis, D.S.P.S.F.G., N.M.F.B; Investigation, D.S.P.S.F.G., N.M.F.B, A.G.O., D.R.R., M.J., J.A.B.S., T.R.A., M.V.C., A.S.V., S.M.H; Resources, A.K., J.R.Z., L.R.S.; Writing – Original Draft, D.S.P.S.F.G., A.K., J.R.Z., L.R.S.; Writing – Review & Editing, D.S.P.S.F.G., A.K., J.R.Z., L.R.S.; Visualization, D.S.P.S.F.G.; Supervision, A.K., J.R.Z., L.R.S.; Project Administration, A.K., J.R.Z., L.R.S.; Funding Acquisition, A.K., J.R.Z., L.R.S.;

## Acknowledgments

We would like to appreciate the technical support of Ann-Marie Petterson. This work was supported by São Paulo Research Foundation (FAPESP) (thematic projects and scholarships: 2016/23008-5, 2017/24795-3, 2021/07411-2, 2018/20581-1, 2017/24851-0), by the Karolinska Institutet Grants with support from Swedish Diabetes Foundation (DIA2021-645), Swedish Research Council for Sport Science (P2019-0140 and P2020-0064), and the Swedish Research Council (2015-00165). This study was financed in part by the Coordenação de Aperfeiçoamento de Pessoal de Nível Superior – Brasil (CAPES) – Finance Code 001 and by the Conselho Nacional de Desenvolvimento Científico e Tecnológico (CNPq).

---

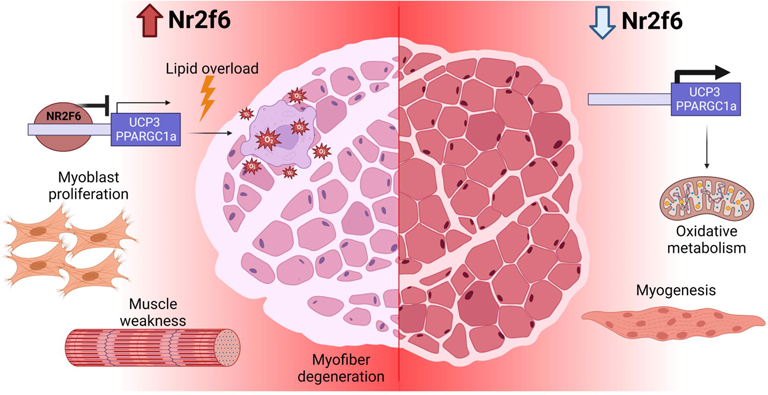

