## Supplementary Figures for "Concerted regulation of skeletal muscle metabolism and contractile properties by the orphan nuclear receptor Nr2f6"

**Contents:**

Supplementary Figures 1 - 5

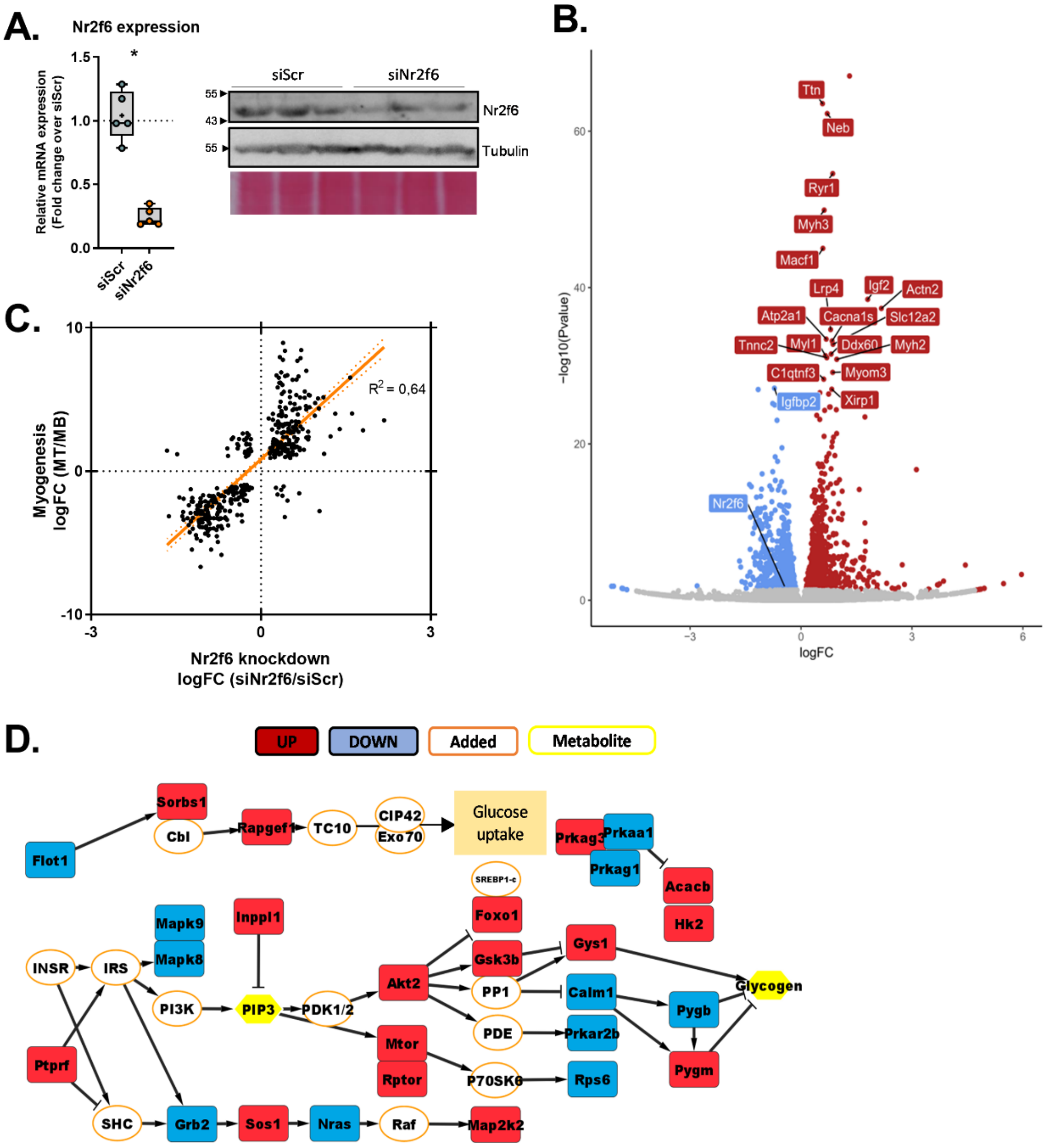


| Up regulated | | | Downregulated | | |
|---|---|---|---|---|---|
| Gene | log2FC | FDR | Gene | log2FC | FDR |
| Sorbs1 | 0.43 | 1.4E-07 | Flot1 | -0.27 | 1.6E-02 |
| Rapgef1 | 0.22 | 9.6E-03 | Sorbs1 | 0.43 | 1.4E-07 |
| Inppl1 | 0.34 | 1.8E-07 | Mapk9 | -0.35 | 8.7E-03 |
| Ptprf | 0.65 | 2.0E-13 | Mapk8 | -0.36 | 1.5E-03 |
| Sos1 | 0.27 | 3.6E-02 | Grb2 | -0.38 | 5.6E-03 |
| Akt2 | 0.29 | 4.9E-05 | Nras | -0.22 | 3.1E-02 |
| Mtor | 0.27 | 4.3E-03 | Prkaa1 | -0.26 | 7.0E-03 |
| Rptor | 0.27 | 1.0E-03 | Prkag1 | -0.27 | 4.5E-02 |
| Map2k2 | 0.24 | 2.7E-02 | Calm1 | -0.24 | 8.4E-05 |
| Foxo1 | 0.56 | 1.3E-04 | Prkar2b | -0.39 | 3.3E-04 |
| Gsk3b | 0.25 | 5.4E-03 | Rps6 | -0.25 | 4.6E-05 |
| Gys1 | 0.45 | 1.6E-10 | Pygb | -0.40 | 4.6E-05 |
| Acacb | 0.78 | 3.2E-03 | | | |
| Hk2 | 0.34 | 1.2E-06 | | | |
| Pygm | 0.64 | 7.0E-17 | | | |

**Fig. S1. Nr2f6 regulates myogenesis and binds to the promoters of genes involved in metabolism in different cell types.** (A) Validation of Nr2f6 knockdown in siScr and siNr2f6 transfected C2C12 myotubes by RT-qPCR (left-most) and western blot (on the right most. (B) Volcano plot of Nr2f6 knockdown C2C12 myocytes. Genes upregulated in red and downregulated in blue (FDR <0.05). N=4-5. (C) Correlation of differentially expressed genes in the transcriptome of siNr2f6 myocytes and public C2C12 differentiation microarray (GSE4694). (D) Manually selected insulin signaling pathway schematic displaying differentially expressed genes after Nr2f6 knockdown and other components of the pathway. Metabolites are depicted in yellow borders and unchanged genes are in orange borders. Fold-change and FDR level of depicted genes in the RNA-seq are shown in the table.

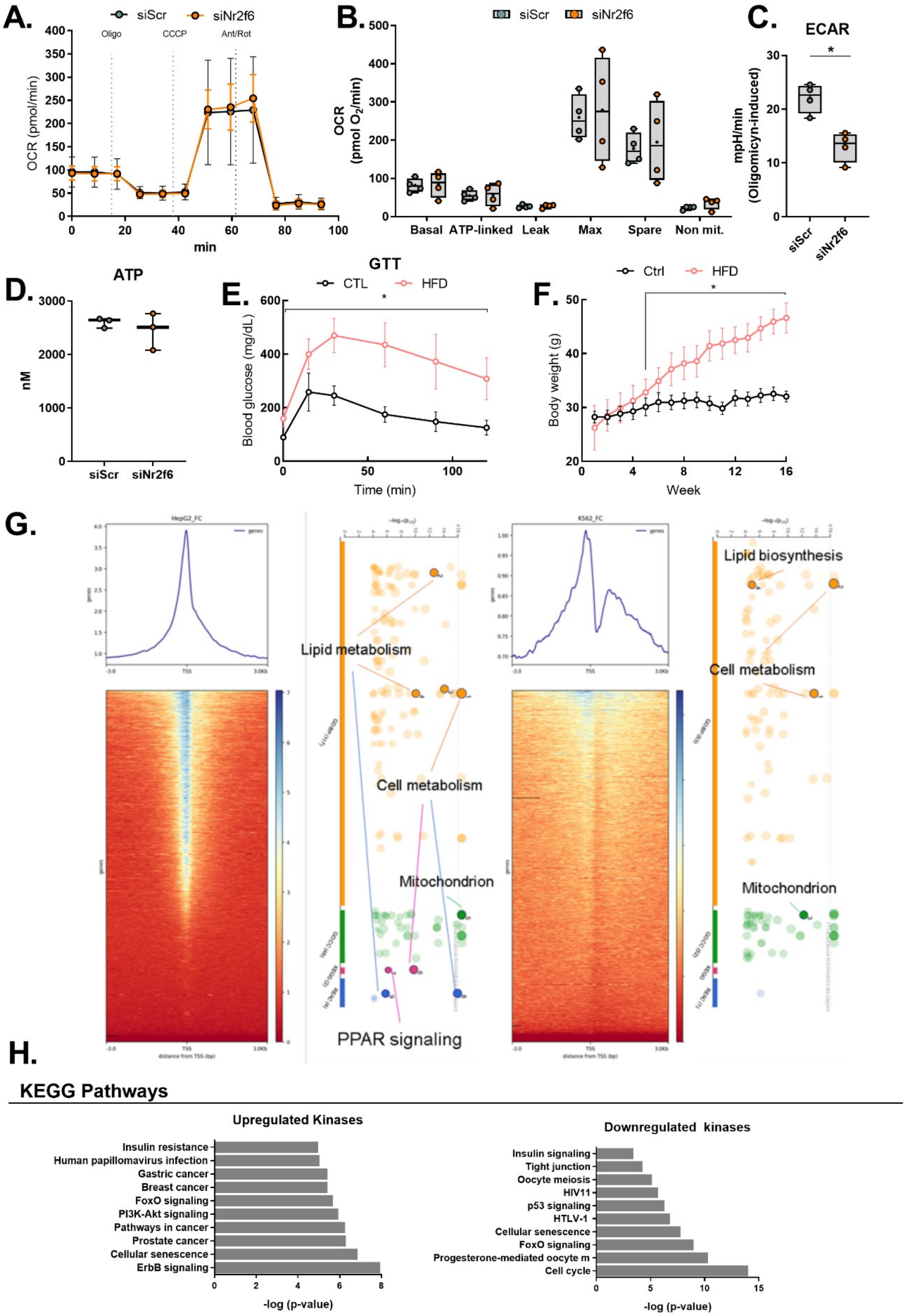
A.
siScr
siNr2f6
Oligo
CCCP
Ant/Rot
OCR (pmol/min)
min
B.
OCR (pmol O2/min)
Basal
ATP-linked
Leak
Max
Spare
Non mit.
C.
ECAR
mpH/min (Oligomicyn-induced)
D.
ATP
nM
E.
GTT
CTL
HFD
Blood glucose (mg/dL)
Time (min)
F.
Ctrl
HFD
Body weight (g)
Week
G.
Lipid metabolism
Cell metabolism
Mitochondrion
PPAR signaling
Lipid biosynthesis
H.
KEGG Pathways
Upregulated Kinases
Insulin resistance
Human papillomavirus infection
Gastric cancer
Breast cancer
FoxO signaling
PI3K-Akt signaling
Pathways in cancer
Prostate cancer
Cellular senescence
ErbB signaling
-log (p-value)
Downregulated kinases
Insulin signaling
Tight junction
Oocyte meiosis
HIV11
p53 signaling
HTLV-1
Cellular senescence
FoxO signaling
Progesterone-mediated oocyte m
Cell cycle
-log (p-value)

**Fig. S2. Nr2f6 depletion enhances metabolism in skeletal muscle.** (A, B) Oxygen consumption assay in C2C12 myocytes transfected with siScr (control) and siNr2f6. On the right, are the calculated metabolic parameters. (C) Oligomycin-induced extracellular acidification rate during a high-glucose oxygen consumption assay. N=4. (D) ATP content in siScr and siNr2f6 myocytes. (E, F) Body weight and glucose tolerance test of mice undergoing 16 weeks of a high-fat diet. (G) Nr2f6 DNA binding profile in the ENCODE ChIP-seq data of HepG2 (right) and K562 (left) cells. Gene ontology analysis of the genes with biding within ±3 kbp of the transcription start site. (H) Enrichment of KEGG pathway terms in the upregulated (left) and downregulated (right) kinases in the siNr2f6 transcriptome. Data displayed as mean ±SD. Boxplot with whiskers spanning minimum to maximal and box edges 25th-75th percentile, the line at the median and + at the mean. * Indicates $p < 0.05$ using unpaired two-tailed Student's t-test.

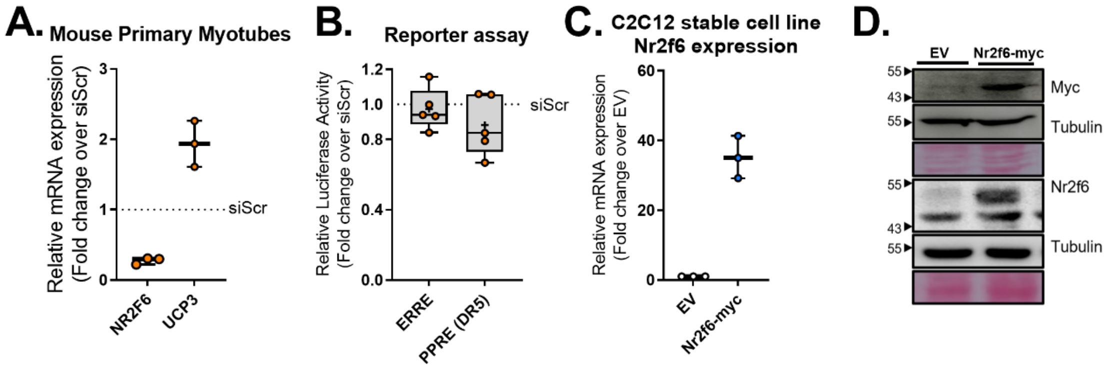


**Fig. S3. Nr2f6 regulates UCP3 and PGC-1α expression.** (A, B) Gene expression (N=3) and representative western blot for validation of Nr2f6-myc stable myotubes. Boxplot with whiskers spanning minimum to maximal and box edges 25th-75th percentile, the line at the median and + at the mean. * Indicates $p < 0.05$ using unpaired two-tailed Student's t-test. (C) Gene expression measured by RT-qPCR in primary mouse skeletal muscle cells transfected with siScr (control) and siNr2f6. N=3. (D) Luciferase reporter assay of the responsive elements of the Estrogen Related Receptor (ERRE) and PPAR (PPRE) in MEF cells transfected with siScr or siNr2f6. N=5.

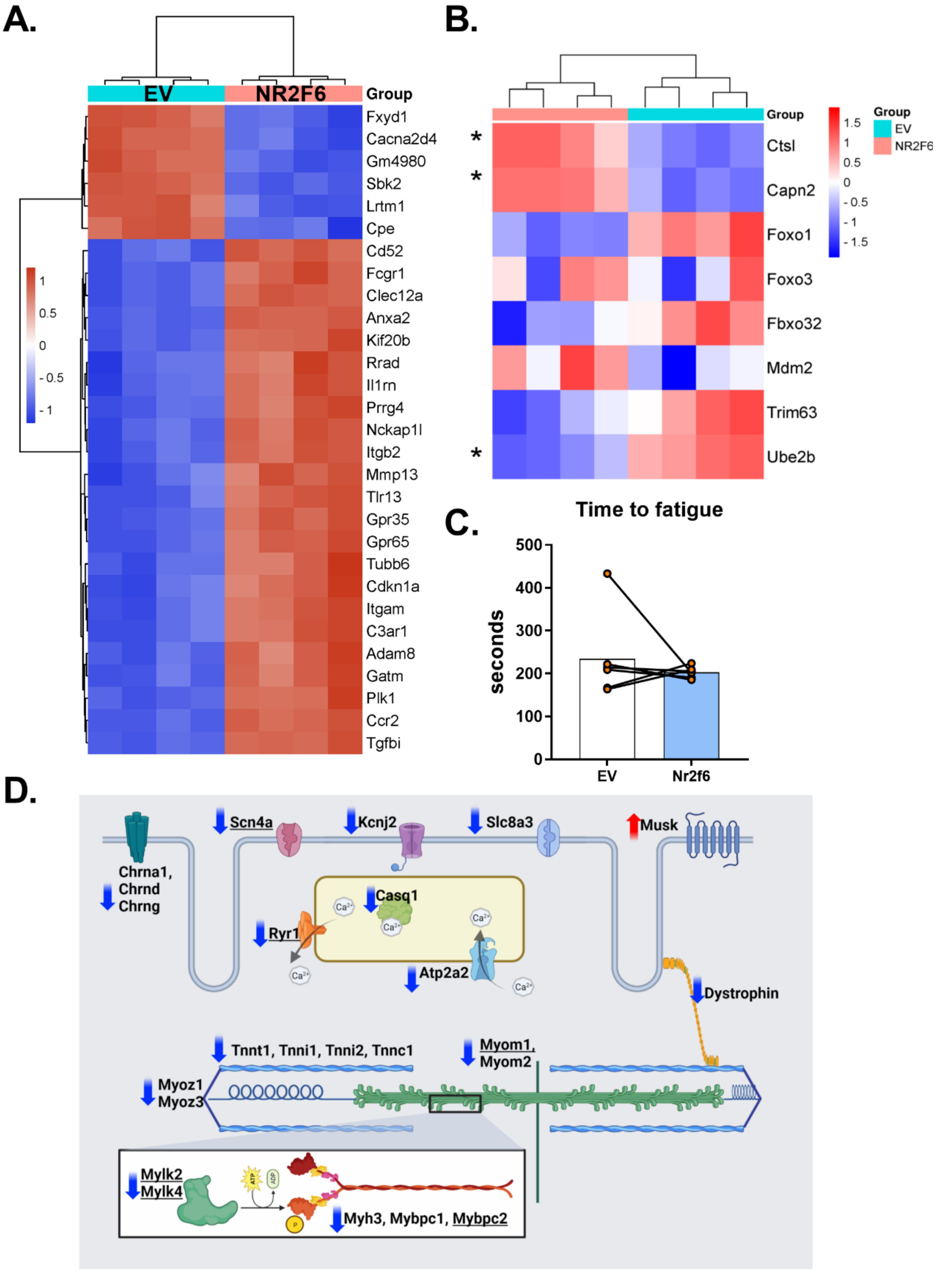


**Fig. S4. Nr2f6 overexpression impairs muscle function.** (A) Heat-map of top 30 most modulated genes in tibialis anterior muscle electroporated with control empty vector (EV) or an Nr2f6-myc-coding plasmid. N=4.(B) Atrogenes regulated by Nr2f6 overexpression in the tibialis anterior muscle. * Denotes significant modulation (FDR < 0.05, fold-change > 2) in the microarray. (C) Time to fatigue in ex vivo contraction was set as the necessary time to reach 50% of the maximal force with constant stimulation. N=6. (D) Nr2f6 overexpression reduces the expression of several genes of the contractile apparatus, myofiber calcium handling, and action potential transduction. Genes with Nr2f6 binding motif at the promoter are underscored. Differentially expressed genes following Nr2f6 overexpression in mouse TA were selected according to ontology terms related to muscle contraction and function. The arrows indicate the up- or downregulation. Sodium Voltage-Gated Channel Alpha Subunit 4 (Scn4a), Potassium Inwardly Rectifying Channel Subfamily J Member 2 (Kcnj2), Solute Carrier Family 8 Member A3 (Slc8a3), Muscle Associated Receptor Tyrosine Kinase (Musk), Ryanodine Receptor 1 (Ryr1), Calsequestrin 1 (Casq1), ATPase Sarcoplasmic/Endoplasmic Reticulum Ca2+ Transporting 2 (SERCA2, Atp2a2), Cholinergic Receptor Nicotinic Alpha 1/delta/gamma subunit Chrna1/d/g), Troponin T1/I1/I2/C1 (Tnnt1/Tnni1/Tnni2/c1), Myom1/2 (Myomesin1/2), Myozenin1/3 (Myoz1/3), Myosin light chain kinase 2/4 (Mylk2/4), Myosin heavy chain 3 (Myh3), Myosin binding protein C1/2 (Mybpc1/2)

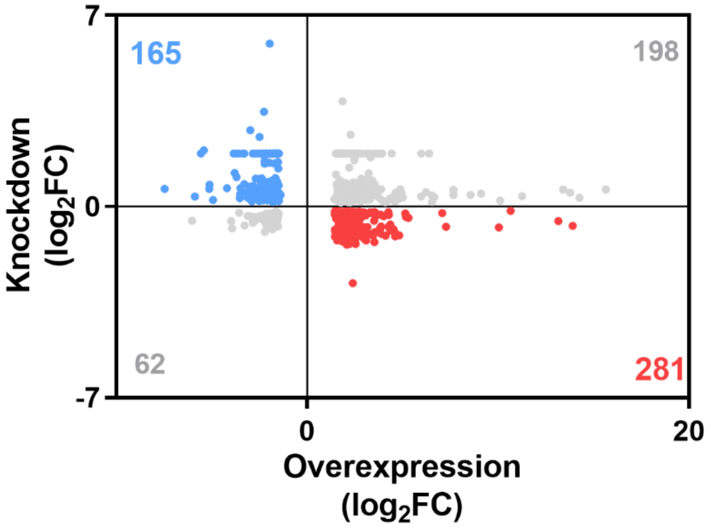


**Fig. S5.** Scatter plot of differentially expressed genes in Nr2f6 knockdown in C2C12 myocytes RNA-seq (FDR < 0.05) and Nr2f6 overexpression in mice TA microarray (FDR <0.05, fold-change >2). In red: genes upregulated by Nr2f6; in blue: genes downregulated by Nr2f6; in grey: genes with the same direction of modulation by Nr2f6 overexpression and knockdown.
